# Contrasting Range and Demographic Histories in two Palearctic Dragonfly Species (Odonata: Anisoptera)

**DOI:** 10.64898/2026.11.05.764621

**Authors:** Aaron M. Goodman, Ethan R. Tolman, Manpreet K. Kohli

**Author notes:** Joint first authors.

## Abstract

Climatic suitability is often used to infer historical range changes, yet whether suitable area predicts effective population size remains unexplored. We investigated this relationship in two Palearctic dragonflies with contrasting ecologies: *Somatochlora arctica*, a widespread peatland specialist, and *S. meridionalis*, a southern European species associated with shaded streams. We combined demographic reconstructions from individual reference genomes using the pairwise sequentially Markovian coalescent (PSMC), maximum entropy (MaxEnt) species distribution models projected across six paleoclimatic intervals spanning approximately 3.3 million years, and ancestral state reconstructions of peatland specialization. Predicted suitable area was consistently greater for *S. arctica*, reaching its greatest extent during the Last Glacial Maximum. However, its inferred effective population size was smaller than that of *S. meridionalis* for most of the last ∼275,000 years. Ancestral reconstructions placed the most probable gain of peatland specialization in the *S. arctica* lineage (Clade I) during the late Miocene, with a median age of 9.4 million years, predating the modeled climatic intervals. These findings indicate a decoupling of climatic suitability and demographic history. Patchy breeding habitat, population structure, and the isolated Scottish origin of the sampled *S. arctica* may contribute to this mismatch. Combining genomic, ecological, and phylogenetic approaches provides complementary insights into historical biogeography and highlights the limitations of interpreting suitable climate as available breeding habitat or using individual genomes to characterize species-wide demographic histories.

## INTRODUCTION

Neutral theory predicts that larger populations harbor greater genetic diversity. In empirical systems, however, genetic diversity is often only weakly correlated with range size and abundance (Lewontin 1974; Leffler et al. 2012; Romiguier et al. 2014). Does the amount of suitable habitat available over time predict long-term trends in effective population size (N_e_)?

### Quaternary Climate as a Natural Experiment

The Quaternary Period (2.58 Ma to present) provides a natural experiment for testing this question. During glacial periods in the Holarctic, temperate species are expected to have retreated into southern refugia, whereas cold-adapted species are expected to have expanded their ranges (Hewitt 1999, 2000, 2004; Stewart et al. 2010). Thus, if N_e_ tracks the availability of suitable habitat, cold-adapted species should show increases in N_e_ during glacial periods, when more suitable habitat is available, and decreases during warmer periods, when less is available. The opposite relationship is expected in warm-adapted species.

### Modeling Historical Biogeography

Several tools can be used to model historical habitat suitability and historical N_e_, and each has major limitations. Species distribution model (SDM) hindcasts relate the current occurrences of a species to climatic variables and project this relationship onto paleoclimate reconstructions to predict where suitable climate may have existed in the past (Nogués-Bravo 2009; Svenning et al. 2011). Such models assume niche conservatism (the stasis of ecological traits over time; Wiens & Graham 2005; Wiens et al. 2010), which may not be realistic in some lineages. Because they are typically built from climatic variables alone, SDM hindcasts also cannot account for the availability of specific habitats, and they provide no direct information on population size.

Demographic models built from population-genomic data sampled across the range of a species (e.g., dadi, Gutenkunst et al. 2009; fastsimcoal2, Excoffier et al. 2013; moments, Jouganous et al. 2017; GADMA2, Noskova et al. 2023; and approximate Bayesian computation [ABC], Beaumont et al. 2002, as implemented in DIYABC Random Forest, Collin et al. 2021) can provide insights into historical population dynamics, including gene flow and changes in population size. Generating such datasets is often prohibitively expensive, and it can be infeasible where behavior and/or regulatory constraints hinder the collection of tissue. Coalescent modeling of a single diploid genome, typically with the pairwise sequentially Markovian coalescent (PSMC; Li & Durbin 2011; Nadachowska-Brzyska et al. 2016), is less detailed than models built from multiple genomes, but it is substantially less expensive. As reference genomes become available for more species (Lewin et al. 2018; Darwin Tree of Life Project Consortium 2022; Mc Cartney et al. 2024), such models can be developed without any new sequencing, provided that the raw reads are publicly available. However, the PSMC does not, on its own, provide evidence of the causes of population expansions or declines, and inferred changes in N_e_ can reflect changes in population structure as well as in population size (Mazet et al. 2016).

Although SDMs and the PSMC could together test the hypothesis of a historical relationship between suitable habitat and N_e_, these tools have rarely been combined (but see Kozma et al. 2018). Here, we combine both tools to investigate the relationship between historical N_e_ and suitable habitat in two Palearctic dragonfly species of the genus *Somatochlora*, a group proposed as an indicator of climate change (Hassall 2015). The first of these species, *S. arctica*, is a specialist of peatlands, and it inhabits colder northern latitudes across Europe and Asia (Boudot & Kalkman 2015; Cristofaro et al. 2024). The second, *S. meridionalis*, is not a peatland specialist and is found in central and southern Europe (Boudot & Kalkman 2015).

Our first objective is to determine whether historical habitat suitability is tied to historical N_e_ in each of these species. If N_e_ tracks suitable habitat, we expect the more widespread, cold-adapted *S. arctica* to have had a larger N_e_ than *S. meridionalis*, with increases during glacial periods. Our second objective is to use ancestral state reconstruction (ASR; Pagel 1994; Huelsenbeck et al. 2003) to determine whether peatland specialization predates the modeled time windows, and whether this specialization can explain any gaps between suitability and N_e_.

We predict that, combined, these three analyses can provide a more holistic view of the biogeography of this system than any one of them could alone. Because it is highly relevant to our findings, we briefly consider the evolution and ecology of the genus *Somatochlora*, including our two focal species, below.

### Evolution and Ecology of Somatochlora

The striped emeralds (Corduliidae: *Somatochlora*) are a Holarctic genus whose larvae (nymphs) occupy habitats ranging from peatlands and seepages to lakes and forest streams (Goodman et al. 2025). Phylogenomic analyses indicate a North American origin during the Miocene, followed by repeated dispersal into Eurasia across Beringia, establishing the genus’ northern distribution prior to Quarternary climatic oscillations (Goodman et al. 2025). Within the Palearctic, species differ widely in their breeding habitats. Nymphal *S. arctica* inhabit small, *Sphagnum*-filled pools, with recent work identifying high moss cover and proximity to woodland as important habitat characteristics (Chrostofaro et al. 2024). In contrast, *S. meridionalis* breeds in shaded streams and small rivers, including watercourses that fragment into pools during the summer (Boudot and Kalkman et al. 2015). These differences suggest that changes in climatic suitability may result in different demographic responses depending on availability and continuity of breeding habitat. Furthermore, an already established time-divergence framework for *Somatochlora* (Goodman et al. 2025) allows us to investigate when peatland specialization arose, providing historical context for interpreting the relationship between suitable habitat and demographic change in *S. arctica*.

## METHODS

### Demographic Modeling

#### Specimens and data

We used publicly available genome assemblies of an adult male *S. meridionalis* sampled from Strandzha Nature Park, Bulgaria (primary assembly, European Reference Genome Atlas; Mc Cartney et al. 2024), and of an adult *S. arctica* collected from Beinn Eighe National Nature Reserve, Scotland (haplotype 1 of the Darwin Tree of Life assembly; Darwin Tree of Life Project Consortium 2022), which we inferred to be male from X-chromosome depth (Table S1). Both genomes were sequenced with PacBio HiFi reads, which we downloaded from the NCBI Sequence Read Archive with the SRA Toolkit (Katz et al. 2022); assemblies were downloaded with NCBI Datasets (O’Leary et al. 2024). For *S. arctica*, we pooled the two HiFi runs from the same BioSample (∼41× raw coverage; median autosomal depth 48.5×), whereas the single HiFi run for *S. meridionalis* provided ∼28× raw coverage (median autosomal depth 28.2×; Table S1).

#### Implementing the PSMC

To model changes in historical effective population size (*N_e_*), we followed an established pipeline (Tolman et al. 2023,[CL1] 2025a,b) for applying the pairwise sequentially Markovian coalescent (PSMC; Li & Durbin 2011) to Odonata genomes assembled with PacBio HiFi data (see supplemental note S1 for details). Heterozygosity was summarized from the PSMC input files (supplemental note S1), in which fq2psmcfa divides each chromosome into 100-bp bins and codes each bin as heterozygous (at least one heterozygous site), homozygous, or missing (more than 90% of the bin’s bases filtered out). We report the fraction of non-missing bins that are heterozygous (heterozygous bins), which approximates 100 times the per-site heterozygosity and is a lower bound because a bin with several heterozygous sites is counted once (Tables S2, S3). Per-chromosome values, expressed relative to the median of the autosomes, were combined with relative depth to identify sex chromosomes and other anomalous chromosomes (Table S2).

### Species Distribution Modeling

#### Occurrence and Environmental Data

We obtained occurrence records for *S. arctica* and *S. meridionalis* from Kalkman et al. (2022) and supplemented them with GBIF records possessing verified coordinates, photographs, observation dates, and at least two-thirds agreement on species identification. We removed duplicate coordinates and spatially thinned records by 5 km using *GeoThinneR* (Mestre-Tomás 2025) to reduce sampling bias and clustering (Veloz 2009).

We acquired 19 CHELSA bioclimatic variables averaged across 1979–2013 at 2.5 arc-minute resolution in WGS84 (Karger et al. 2023). We excluded bio08 and bio09 because of spatial artifacts (Booth 2022) and bio02, bio03, bio05, bio06, and bio07 because they were unavailable in our paleoclimate datasets (Brown et al. 2018). Rasters were cropped to 25°W– 180°E and 15–85°N (Palearctic spatial extent, Olson et al. 2001), and correlated predictors were removed using *usdm* at an absolute Pearson correlation threshold of 0.7 (Naimi et al. 2025; Goodman et al. 2023). All analyses were conducted in R v4.6.1 (Team 2021).

We defined species-specific study extents using 200-km buffers around occurrences (Peterson and Soberón 2012). To account for sampling bias, we generated a kernel-density bias raster from all odonate records in Kalkman et al. (2022) using *ks* (Duong 2007; Phillips et al. 2009; Kramer-Schadt et al. 2013). Within each calibration extent, we sampled 50,000 background points with probabilities proportional to bias-raster values.

#### Model Building

We built Maxent presence-background models (Phillips et al. 2017; Phillips 2021) using *ENMeval* (Kass et al. 2021) and evaluated them with four-region spatial block cross-validation, training on three regions and withholding the fourth (Roberts et al. 2017; Valavi et al. 2021). We tuned complexity using four feature-class combinations (L, LQ, Q, and LQH) and regularization multipliers of 1–4, yielding 16 candidate models per species (Warren and Seifert 2011; Radosavljevic and Anderson 2014).

We assessed models using mean validation Area Under the Curve (AUC), 10-percentile omission rate, and corrected Akaike Information Criterion (AICc) calculated from the full calibration dataset. Among models with ΔAICc < 2, we selected the model with the lowest mean omission rate, resolving ties by highest mean validation AUC, followed by lowest AICc. Final models were fitted to the full calibration dataset.

We examined response curves by varying each predictor while holding the others at their model sample means (Phillips et al. 2017). We calculated permutation importance by randomly shuffling each predictor among presence and background points and measuring the resulting decrease in training AUC (Phillips 2021). Positive AUC decreases were normalized to percentages to compare relative importance among predictors.

#### Suitability Predictions

We generated present-day suitability predictions across our Palearctic extent using the complementary log-log transformation (Phillips 2021) and converted them to binary maps using the 10th percentile of suitability at retained occurrence localities. We projected selected models onto PaleoClim variables (Brown et al. 2018), applying the same species-specific thresholds, for the mid-Holocene (8.326–4.2 ka), Last Glacial Maximum (∼21 ka), Last Interglacial (∼130 ka), MIS19 (∼787 ka), and MIS M2 (∼3.3 Ma).

#### Comparison of Suitable Habitat and Demographic Histories

We calculated suitable area by summing cell areas in binary predictions using *terra* (Hijmans et al. 2022). We excluded ice-covered areas using ICE-6G_C reconstructions at 6 ka for the mid-Holocene and 21 ka for the LGM (Argus et al. 2014; Peltier et al. 2015), and the 130- ka reconstruction of Menviel et al. (2019) for the Last Interglacial. The 6-ka reconstruction served as a proxy for the mid-Holocene interval; present-day, MIS19, and MIS M2 predictions remained unmasked.

### Ancestral State Reconstruction of peatland Habitat Specialization

We fit continuous-time Markov (Mk) models for discrete characters (Pagel 1994; Lewis 2001) by maximum likelihood with fitMk in phytools (Revell 2012, 2024) to assess whether the *S. arctica* lineage is likely to have been a peatland specialist throughout the period spanned by the PSMC and niche models (habitat codings in Table S4). To estimate the number of gains and losses of peatland specialization across the tree, and when the *S. arctica* lineage most recently gained this state, we used stochastic character maps (Nielsen 2002; Huelsenbeck et al. 2003; Bollback 2006) simulated with make.simmap (phytools; see supplemental note S2 for detailed methodology).

## RESULTS

### Demographic modeling

More than 99.6% of primary HiFi reads mapped to their respective assemblies (Table S1). We identified the unlabelled X chromosome of *S. meridionalis* as chromosome 12 (OZ317816.1), which had 0.51× the autosomal depth and 0.07× the median autosomal heterozygosity; the labelled X of *S. arctica* (OZ407525.1) had 0.50× the autosomal depth, consistent with an XO male (Table S2). Approximately 34% of the non-missing 100-bp bins of *S. meridionalis* were heterozygous, compared with 18% for *S. arctica* (chromosome 5 inversion masked and microchromosome excluded; Table S3). Correspondingly, the population-scaled mutation rate estimated by PSMC (θ_₀_ = 4N_₀_μ per 100-bp bin, which reflects the average divergence between the two haplotypes of each genome) was about 2.5 times higher in *S. meridionalis* (0.277) than in *S. arctica* (0.113). Under our mutation rate, these values correspond to reference effective population sizes (N_₀_, the size against which PSMC expresses population size in each time interval) of 346,708 and 141,134, respectively (Table S3).

All three *S. arctica* PSMC models produced consistent trajectories (Fig. S1; Table S3). Masking the chromosome 5 inversion and excluding the microchromosome mainly lowered the ancient peak (from 1.37 to 1.25 million), and additionally excluding chromosome 9 lowered the Late Pleistocene minimum (from 10,600 to 8,500). We therefore present the model with the inversion masked and the microchromosome excluded in Fig. 1 (top). Both species show an ancient peak in *N_e_* (*S. meridionalis*, 1.34 million at 10.6–11.7 Ma; *S. arctica*, 1.25 million at 7.6– 8.4 Ma) and a smaller, more recent peak (*S. arctica*, 732,000 at 541–603 ka; *S. meridionalis*, 588,000 at 729–821 ka; Fig. 1, top). Late Pleistocene minimum Ne differed approximately seven-fold between the species (*S. arctica*, 10,600 at 12–19 ka; *S. meridionalis*, 71,500 at 21– 43 ka; Fig. 1, top).

**Figure 1.**
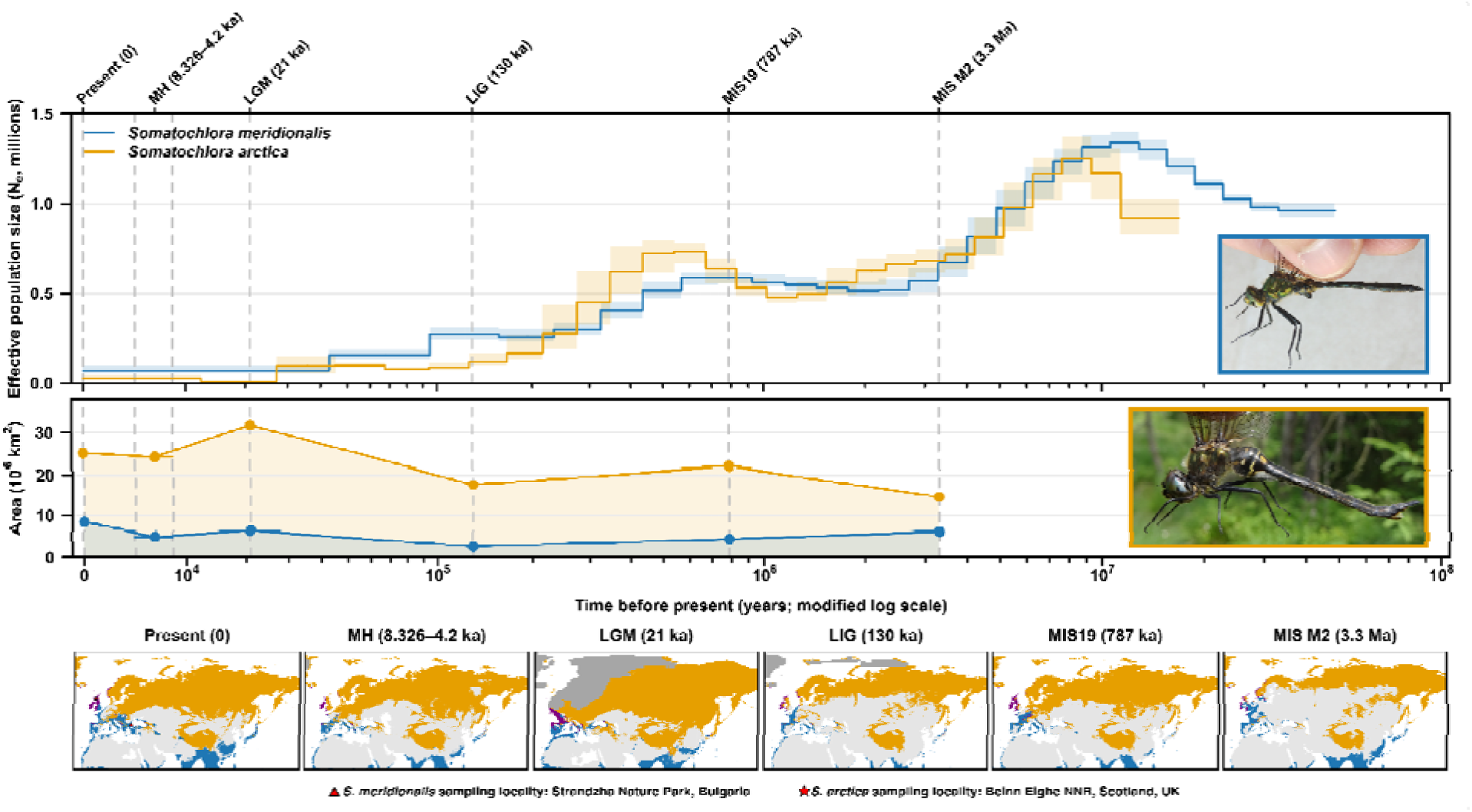
Demographic histories and predicted climatic suitability of *Somatochlora arctica* (orange) and *S. meridionalis* (blue). Top: effective population size (N_e_) inferred using PSMC, with shaded envelopes showing bootstrap variation. Middle: total predicted suitable area calculated from thresholded Maxent predictions for six climatic intervals. Dashed vertical lines mark the present, mid-Holocene (MH), Last Glacial Maximum (LGM), Last Interglacial (LIG), MIS19, and MIS M2. Bottom: Corresponding predictions of suitable habitat across the Palearctic; purple indicates overlap between species, and gray indicates ice-covered areas. Ice-covered areas were excluded from MH, LGM, and LIG projections. Photographs: *S. arctica*, © Christoph Moning (iNaturalist photo 60305072; CC BY 4.0); *S. meridionalis*, Harukano (iNaturalist photo 406561275; CC0). Photographs cropped to fit.

### Species Distribution Modeling

Both optimal models included linear, quadratic, and hinge feature classes, with regularization multipliers of four for *S. arctica* and two for *S. meridionalis*. Mean validation AUC was high for *S. arctica* (0.85) and adequate for *S. meridionalis* (0.76), while 10-percentile omission exceeded its expected value of 0.1 (0.14 and 0.24, respectively). Optimal models possessed 23 and 24 non-zero coefficients, respectively.

Mean temperature of the warmest quarter (bio10) possessed the highest permutation importance for both species (72.51% for *S. arctica*; 60.00% for *S. meridionalis*), followed by temperature seasonality (bio04) for *S. arctica* (15.17%) and precipitation of the coldest quarter (bio19) for *S. meridionalis* (16.02%). Bio10 expressed asymmetric quadratic-like relationships, with greatest suitability at 13–15°C and 25°C, respectively. Precipitation seasonality (bio15) expressed jagged quadratic-like relationships, with optima near 30% and 51%, while precipitation of the warmest quarter (bio18) produced greatest suitability at 335–410 mm and 410 mm, respectively. Precipitation of the coldest quarter (bio19) expressed a positive, saturating relationship for *S. meridionalis* above approximately 420 mm (Figs S2–S5).

Present-day predictions revealed suitable habitat across northern Eurasia for *S. arctica*, with additional areas within central European mountains and the Tibetan Plateau. Suitability for *S. meridionalis* encompassed western and southern Europe, the Mediterranean, and portions of southern Asia (Fig. 1, bottom; Figs S6, S7).

For *S. arctica*, suitable area expanded southward during the LGM before contracting during the mid-Holocene. For *S. meridionalis*, suitable area increased from the Last Interglacial to the LGM, contracted during the mid-Holocene, and expanded toward the present. Suitable area remained greater for *S. arctica* across all modeled intervals (Fig. 1, middle and bottom; Figs S6, S7).

### Ancestral State Reconstruction of Peatland Habitat Specialization

Under our primary coding, ER was marginally preferred (AICc = 48.36, weight = 0.54) over ARD (ΔAICc = 0.34, weight = 0.46; Table 1). ER was also marginally preferred under all three alternative codings (weights 0.55–0.68; Table 1). Under the ARD model fitted to the primary coding, the loss rate exceeded the gain rate (q(loss) = 0.092 vs. q(gain) = 0.026 per lineage per Myr; Table 1).

**Table 1.** Mk models fitted to peatland specialization in *Somatochlora* (40 species) under the primary and alternative codings (numbered as in the Methods).

| **Coding** | **Model** | **k** | **log L** | **AICc** | **ΔAICc** | **Weight** | **q gain (Myr<sup>−1</sup>)** | **q loss (Myr<sup>−1</sup>)** |
|---|---|---|---|---|---|---|---|---|
| Primary coding | ER | 1 | -23.13 | 48.36 | 0.00 | 0.54 | 0.027 | 0.027 |
|  | ARD | 2 | -22.19 | 48.70 | 0.34 | 0.46 | 0.026 | 0.092 |
| Alternative coding 1 | ER | 1 | -22.63 | 47.37 | 0.00 | 0.59 | 0.026 | 0.026 |
|  | ARD | 2 | -21.89 | 48.11 | 0.74 | 0.41 | 0.023 | 0.084 |
| Alternative coding 2 | ER | 1 | -25.65 | 53.41 | 0.00 | 0.68 | 0.046 | 0.046 |
|  | ARD | 2 | -25.32 | 54.96 | 1.55 | 0.32 | 0.043 | 0.071 |
| Alternative coding 3 | ER | 1 | -19.95 | 42.01 | 0.00 | 0.55 | 0.017 | 0.017 |
|  | ARD | 2 | -19.06 | 42.44 | 0.43 | 0.45 | 0.015 | 0.096 |

Across stochastic maps, peatland specialization was both gained (mean 7.5; 95% interval 3–11) and lost (mean 6.1; 95% interval 1–17) independently. The crown ancestral state (23.7 Ma) was equivocal, but was most likely not a peatland specialist (P(specialist) = 0.29; Fig. 2a). Ancestors with high specialist probability occurred mainly in the clade containing *S. arctica*: the *S. arctica*–*S. incurvata* crown node (5.7 Ma) had P = 0.93, and the *S. whitehousei* + *S. septentrionalis* ancestor had P = 0.97 (Fig. 2a). In 85% of maps, the most recent gain of peatland specialization on the *S. arctica* lineage fell within the crown. The most probable placement of this gain was the stem of the *S. arctica* + *S. franklini* + *S. forcipata* + *S. dido* + *S. incurvata* clade (Clade I, Goodman et al. 2025; branch e; 13.7–5.7 Ma; 64% of maps; median 9.4 Ma, late Miocene; Fig. 2b; Table S5). Across maps with a gain inside the crown, the median gain age was 9.6 Ma (95% HDI 0.2–17.1 Ma). Under every coding, branch e was the single most probable placement of the most recent gain (36–68% of maps; median age 9.3–9.6 Ma; Fig. 2b–e). Reconstructions under the alternative codings were largely consistent with the primary coding (Figs S8–S10), although alternative coding 2 shifted probability toward older branches (branches a–d together 40% of maps vs. 11% under the primary coding).

**Figure 2.**
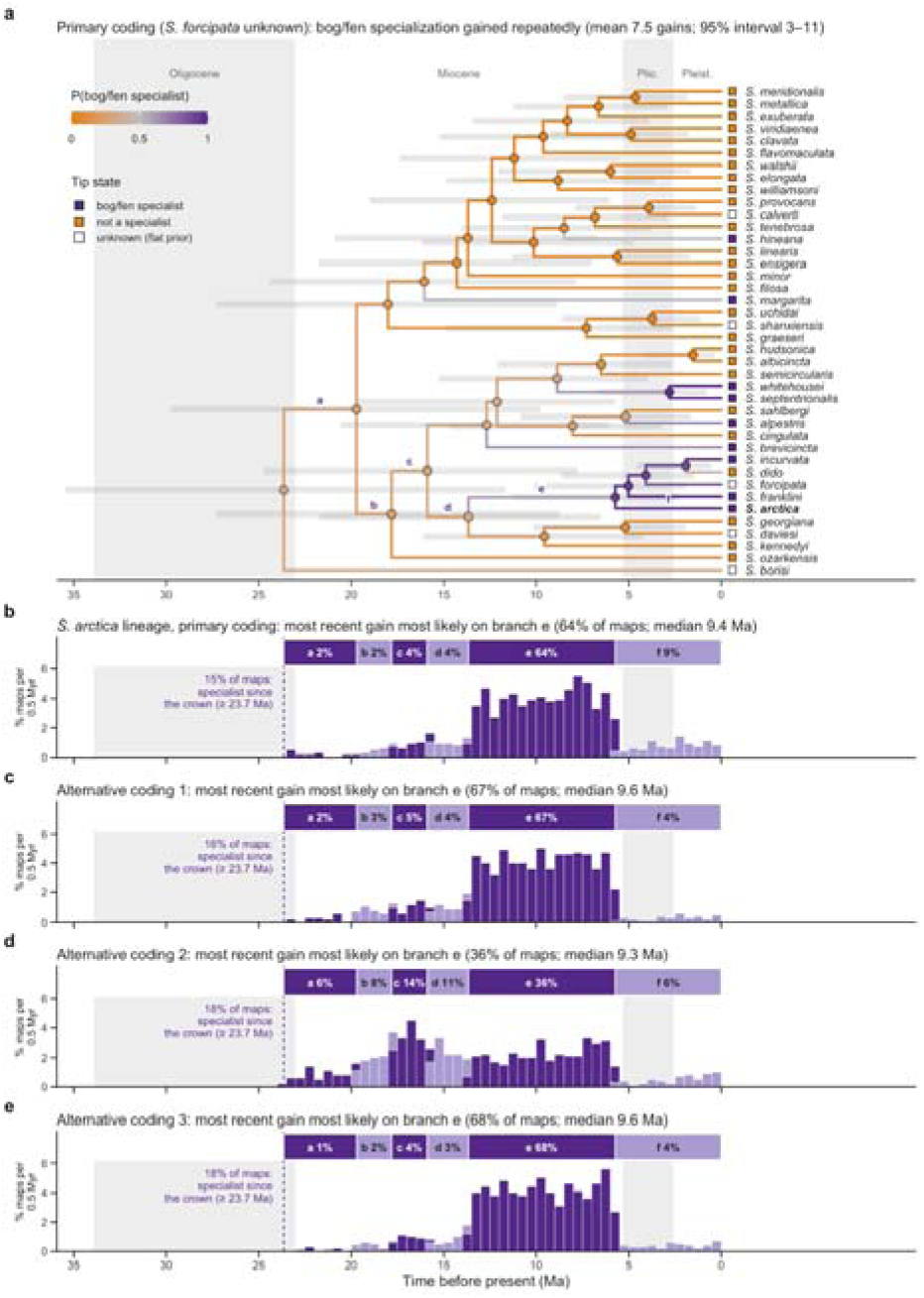
Evolution of peatland specialization in *Somatochlora* and timing of its most recent gain on the *S. arctica* lineage. (a) Time-calibrated tree of 40 species, primary coding (Table S4, with *S. forcipata* treated as unknown). Branch color is the posterior proportion of each branch in the peatland state (1,000 model-averaged stochastic maps). Node symbols are model-averaged marginal ancestral-state probabilities. Gray bars are 95% HPD node ages. Tip squares give the coded state (open: unknown, flat prior). Letters a–f mark the branches from the crown to *S. arctica* (bold). Shading marks geological epochs (Cohen et al. 2013). (b) Posterior age of the most recent peatland gain on the *S. arctica* lineage (start of the uninterrupted peatland run ending at *S. arctica*), in 0.5-Myr bins colored by branch. The strip above each histogram shows the span of branches a–f and the percentage of maps placing the gain on each. Maps in which the lineage was specialist throughout (gain at or before the crown; dashed line) are given as a percentage and not plotted. (c–e) As (b), under alternative codings 1–3: *S. forcipata* coded as a specialist (c); *S. forcipata* and four conservative ‘No’ codings coded ‘Yes’ (d); and two qualified ‘Yes’ codings recoded ‘No’ (e). Panels b–e share the y-axis scale.

## DISCUSSION

Although caution is warranted in tying shifts in historical N_e_ to specific climatic events, especially because generation times and genome-wide mutation rates influence the inferred timing of coalescent events (Tolman et al. 2023; Tolman et al. 2025), we generally find a decoupling of climatically suitable area and N_e_. *S. arctica* is predicted to have had more suitable area in the recent past (20 ka to present) than in the deeper past (>100 ka; Fig. 1, middle), yet the PSMC infers a smaller N_e_ in the most recent atomic intervals and a much larger N_e_ deeper in time (Fig. 1, top). The suitable area of *S. meridionalis* is largely static (Fig. 1, middle), whereas the PSMC infers a much higher N_e_ deeper in time (Fig. 1, top). Moreover, although *S. arctica* is predicted to have had several times more suitable area than *S. meridionalis* in every time slice, its N_e_ has not been consistently larger: for most of the last ∼275,000 years it has been smaller (Fig. 1, top), and its estimated heterozygosity is roughly half that of *S. meridionalis* (Table S3).

### Integrative Methods Provide a Clearer Picture of the Past

Interpreted on their own, each of our three analyses (PSMC, SDM, ASR) could lead to erroneous or incomplete conclusions in this system. On its own, the PSMC could be interpreted as evidence of a widespread decline in the N_e_ of *S. arctica* (Fig. 1, top). The SDMs alone would suggest that *S. arctica* has always had larger populations than *S. meridionalis*, with a large expansion at the LGM (Fig. 1, middle), as expected for a cold-adapted species (Stewart et al. 2010), whereas the ASR alone would provide an approximate time period for specialization without any clear demographic consequences (Fig. 2). Combined, the PSMC and SDMs suggest that suitable climate, as modeled with the available variables, is not an effective proxy for N_e_, consistent with the weak relationship between range size or abundance and genetic diversity across animals (Leffler et al. 2012; Romiguier et al. 2014). The SDMs, which predict unsuitable climate for *S. arctica* in northern France (Fig. 1, bottom), provide context for the very low recent N_e_ of the sampled *S. arctica* (approximately 10,600 at 12–19 ka; Fig. 1, top). This low N_e_ could reflect the isolation of populations in the British Isles, resulting either from colonization or from the loss of connecting habitat. Regardless of the scenario, *S. arctica* must have colonized Scotland relatively recently, because the sampled locality lay beneath the British–Irish Ice Sheet at the LGM (Clark et al. 2022). A recent founder event (Excoffier et al. 2009) is consistent with the low recent N_e_ inferred by the PSMC, although the PSMC has limited resolution in the most recent intervals (Li & Durbin 2011).

Key to both the historical SDMs and our interpretation of the PSMC is the assumption that historical populations had the same habitat requirements as present populations. For example, could past decreases in the N_e_ of *S. arctica* be tied to its colonization of peatland habitats? Our ASR suggests that this is unlikely: the most recent gain of peatland specialization on the *S. arctica* lineage most probably occurred before 5.7 Ma (median 9.4 Ma; Fig. 2), although a more recent gain, on the terminal *S. arctica* branch, remains a small possibility (4– 9% of stochastic maps across habitat codings; Table 2). *S. arctica* has therefore most likely been a peatland specialist throughout the period covered by our SDMs and most, if not all, of the period covered by the PSMC.

**Table 2.** Sensitivity of the reconstruction to habitat coding. Branch columns give the percentage of 1,000 maps placing the most recent gain on the *S. arctica* lineage on each branch (letters as in Fig. 2a) or before the crown. Alternative codings are numbered as in the Methods.

| Coding | Gains (95% int.) | P(crown) | Before crown | a | b | c | d | e | f | Median e (Ma) |
|---|---|---|---|---|---|---|---|---|---|---|
| Primary coding | 7.5 (3–11) | 0.29 | 15 | 2 | 2 | 4 | 4 | 64 | 9 | 9.4 |
| Alternative coding 1 | 7.1 (2–11) | 0.27 | 16 | 2 | 3 | 5 | 4 | 67 | 4 | 9.6 |
| Alternative coding 2 | 10.3 (4–16) | 0.32 | 18 | 6 | 8 | 14 | 11 | 36 | 6 | 9.3 |
| Alternative coding 3 | 4.8 (1–8) | 0.30 | 18 | 1 | 2 | 4 | 3 | 68 | 4 | 9.6 |

### Decoupling of N_e_ and suitable climate

Several factors could explain the observed decoupling of N_e_ and predicted climatic suitability. The habitat specialization of *S. arctica* likely plays a major role. Peatlands are patchy, which implies metapopulation dynamics that affect N_e_ in two opposing ways: local extinction and recolonization reduce the long-term N_e_ of a subdivided population (Whitlock & Barton 1997), whereas population structure inflates estimates of N_e_ from the PSMC and can produce an apparent decline toward the present even when total population size is constant (Mazet et al. 2016). Such structure could explain why, before ∼275 ka, *S. arctica* had an N_e_ similar to or larger than that of *S. meridionalis* (Fig. 1, top), a species that is observed at much higher densities than *S. arctica* despite its more limited range. Further structure is likely added because the sampled individual belongs to a highly isolated population. Finally, glacial climates may have been suitable for *S. arctica*, but they provided poor conditions for peat formation, and most northern peatlands formed after the LGM (MacDonald et al. 2006). The large suitable area predicted for *S. arctica* at the LGM (Fig. 1, middle) may therefore greatly overestimate the breeding habitat that was actually available.

*S. meridionalis* has a small but generally stable range (Fig. 1, middle and bottom) centered on the Balkan Peninsula, one of the principal Pleistocene refugia of European biota (Hewitt 1999; Tzedakis et al. 2002; Schmitt 2007). A small range that persists through glacial cycles may sustain a higher long-term N_e_ than a large range that repeatedly expands, contracts and shifts (Hewitt 2000; Carnaval et al. 2009). Our SDMs show suitable climate for this species across parts of North Africa, Europe and Asia (Fig. 1, bottom). However, climatic suitability does not mean that *S. meridionalis* has dispersed, or will disperse, to these areas. Unlike *S. arctica*, it does not seem to occupy most of its climatically suitable area in the Palearctic.

### Conservation implications

Both individual genomes (Brüniche-Olsen et al. 2021; Formenti et al. 2022; Wilder et al. 2023) and SDMs (Guisan et al. 2013; Syfert et al. 2014) have been used to assess the conservation status of species. Here, we demonstrate that these tools can be used in a complementary manner. The comparatively low heterozygosity of the sampled *S. arctica* (18.3% of 100-bp bins heterozygous, compared with 34.0% in *S. meridionalis*; Table S3) is consistent with its listing as Near Threatened in Great Britain, where it is restricted to isolated sites in Scotland (Daguet et al. 2008; Cristofaro et al. 2024). The SDMs suggest that this low diversity may not be representative of the entire range of *S. arctica*, as the reference genome comes from a disjunct population (Fig. 1, bottom). Such information should, of course, be considered alongside other factors. Although they may have higher genetic diversity than populations from the British Isles, populations of *S. arctica* from mainland Europe are nonetheless threatened by the warming and drainage of peatlands (Tanneberger et al. 2017; Kalkman et al. 2018).

## Conclusions

We find here that climatic suitability is a poor proxy for demographic history in *S. meridionalis* and *S. arctica*. Nonetheless, we demonstrate that demographic modeling from reference genomes, species distribution models, and ancestral state reconstruction can together provide complementary insights into historical biogeography in the absence of in-depth population-genomic sampling. As initiatives such as the Earth BioGenome Project, the Darwin Tree of Life and the European Reference Genome Atlas generate reference genomes for thousands of species (Lewin et al. 2018; Darwin Tree of Life Project Consortium 2022; Mc Cartney et al. 2024), and species occurrence data grows through community science initiatives, this integrative approach can be applied to many more taxa.

