## Supplementary material for "Contrasting Range and Demographic Histories in two Palearctic Dragonfly Species (Odonata: Anisoptera)": Tables S1-S5, Supplemental Note S1-S2, Figure S1-S10

**Table S1.** Specimens, reference assemblies and PacBio HiFi data used for PSMC analyses of *S. meridionalis* and *S. arctica*.

|  | *S. meridionalis* | *S. arctica* |
| --- | --- | --- |
| ***Specimen*** | | |
| Specimen ID | ERGA SK 0506 00332 | SAN0001780 |
| Voucher | BG-NMNHS-ENT-000000019725 | Not provided |
| BioSample | SAMEA117079340 | SAMEA12997840 |
| Locality | Strandzha Nature Park, Izgrev, Bulgaria | Beinn Eighe NNR, Highlands, Scotland, UK |
| Coordinates | 42.1356 N, 27.8042 E | 57.6093 N, 5.3132 W |
| Habitat | Creek in deciduous forest | Mixed upland heath and pinewood |
| Collection date | 11 July 2024 | 21 July 2021 |
| Sex | Male^a^ | Male^b^ |
| ***Reference assembly*** | | |
| Assembly | ioSomMeri1.1 | ioSomArct2.hap1.1 |
| Accession | GCA_974644565.1 | GCA_978407325.1 |
| Initiative | ERGA | Darwin Tree of Life |
| Total length (Mb) | 2,424.7 | 2,516.2 |
| Chromosome-scale autosomes (Mb) | 1,972.9^c^ | 1,905.8 |
| Sex chromosome (Mb) | —^c^ | 136.4 |
| Unplaced or unlocalised (%) | 18.6 | 18.8 |
| ***PacBio HiFi sequencing*** | | |
| Run (platform; Gb) | ERR15496052 (Revio; 67.2) | ERR16055458 (Sequel IIe; 26.8)^d^ + ERR16055457 (Revio; 76.5) |
| Reads (n) | 5,684,917 | 11,943,032 |
| Primary reads mapped (%) | 99.67 | 99.99 |
| Median autosomal depth (×) | 28.21 | 48.50 |
| Consensus depth filter (×) | 10–56 | 16–97 |

^a^ Recorded at collection; X chromosome at 0.51× autosomal depth. ^b^ Not recorded; X chromosome at 0.50× autosomal depth and XO according to the assembly curation notes. ^c^ Twelve assembled chromosomes; chromosome 12 (OZ317816.1) is hemizygous (unlabelled X) and is included in this total. ^d^ The run used to build the assembly. Both runs are derived from BioSample SAMEA12997840 and were pooled. Reads mapped with minimap2 to each species' own assembly; depth from mosdepth (MAPQ ≥ 20, 10-kb windows).

**Table S2.** Per-chromosome depth, heterozygosity and inclusion in the PSMC analyses.

| Chr. | Accession | Length (Mb) | Median depth (×) | Depth ratio | Missing (%) | Het. bins | Het. ratio | Status |
| --- | --- | --- | --- | --- | --- | --- | --- | --- |
| ***S. meridionalis*** | | | | | | | | |
| 1 | OZ317805.1 | 199.9 | 28.4 | 1.01 | 2.6 | 0.313 | 0.94 | Included |
| 2 | OZ317806.1 | 196.3 | 28.3 | 1.00 | 2.4 | 0.319 | 0.96 | Included |
| 3 | OZ317807.1 | 197.2 | 28.1 | 0.99 | 2.5 | 0.331 | 1.00 | Included |
| 4 | OZ317808.1 | 170.8 | 28.3 | 1.00 | 2.2 | 0.344 | 1.04 | Included |
| 5 | OZ317809.1 | 166.8 | 28.3 | 1.00 | 2.5 | 0.319 | 0.96 | Included |
| 6 | OZ317810.1 | 166.1 | 28.3 | 1.00 | 2.9 | 0.301 | 0.91 | Included |
| 7 | OZ317811.1 | 161.4 | 28.1 | 1.00 | 2.5 | 0.322 | 0.97 | Included |
| 8 | OZ317812.1 | 154.5 | 27.9 | 0.99 | 2.5 | 0.374 | 1.13 | Included |
| 9 | OZ317813.1 | 152.0 | 28.2 | 1.00 | 2.4 | 0.373 | 1.13 | Included |
| 10 | OZ317814.1 | 144.1 | 27.9 | 0.99 | 3.2 | 0.392 | 1.19 | Included |
| 11 | OZ317815.1 | 135.8 | 28.4 | 1.00 | 3.2 | 0.380 | 1.15 | Included |
| 12 | OZ317816.1 | 128.0 | 14.3 | 0.51 | 12.1 | 0.022 | 0.07 | Included |
| ***S. arctica*** | | | | | | | | |
| 1 | OZ407513.1 | 197.8 | 49.2 | 1.01 | 3.2 | 0.139 | 0.84 | Included |
| 2 | OZ407514.1 | 194.0 | 49.1 | 1.01 | 3.4 | 0.169 | 1.02 | Included |
| 3 | OZ407515.1 | 183.1 | 49.1 | 1.01 | 1.0 | 0.155 | 0.93 | Included |
| 4 | OZ407516.1 | 175.9 | 48.6 | 1.00 | 6.3 | 0.162 | 0.97 | Included |
| 5 | OZ407517.1 | 167.2 | 47.8 | 0.99 | 5.6 | 0.296 | 1.78 | Included; inversion masked^b^ |
| 6 | OZ407518.1 | 165.3 | 48.7 | 1.00 | 3.4 | 0.174 | 1.05 | Included |
| 7 | OZ407519.1 | 162.8 | 48.6 | 1.00 | 3.2 | 0.154 | 0.93 | Included |
| 8 | OZ407520.1 | 159.3 | 48.7 | 1.00 | 1.8 | 0.158 | 0.95 | Included |
| 9 | OZ407521.1 | 152.9 | 47.6 | 0.98 | 4.9 | 0.368 | 2.22 | Included^c^ |
| 10 | OZ407522.1 | 140.8 | 48.5 | 1.00 | 2.5 | 0.163 | 0.98 | Included |
| 11 | OZ407523.1 | 133.7 | 48.3 | 0.99 | 1.9 | 0.186 | 1.12 | Included |
| 12 | OZ407524.1 | 73.1 | 36.4 | 0.75 | 15.1 | 0.249 | 1.50 | Excluded (microchromosome)^d^ |
| X | OZ407525.1 | 136.4 | 24.1 | 0.50 | 10.5 | 0.015 | 0.09 | Excluded (X) |

Depth ratio: median depth relative to the autosomal median. Het. bins: fraction of callable 100-bp bins containing ≥ 1 heterozygous site. Het. ratio: relative to the median of depth-passing autosomes. ^a^ Unlabelled X: half depth and near-zero heterozygosity. ^b^ Heterozygous inversion at 65.88–152.04 Mb (assembly curation notes) set to missing (heterozygous bins 0.337 inside vs 0.251 for the rest of the chromosome); values shown are before masking. ^c^ Excluded in the sensitivity model only (Fig. S1). ^d^ Order and orientation uncertain over 11.82–53.83 Mb (assembly curation notes).

**Table S3.** PSMC input and model summaries (iteration 20).

| Model | Seq. (n) | Callable (Mb) | Missing (%) | Het. bins | θ₀ | ρ₀ | *N*₀ | g (y) | Min. recomb. |
| --- | --- | --- | --- | --- | --- | --- | --- | --- | --- |
| ***S. meridionalis*** | | | | | | | | | |
| All autosomes (Fig. 1) | 11 | 1,796.9 | 2.6 | 0.340 | 0.277366 | 0.040698 | 346,708 | 3 | 388 |
| ***S. arctica*** | | | | | | | | | |
| Chr. 5 inversion masked, chr. 12 excluded (Fig. 1) | 11 | 1,687.4 | 7.9 | 0.183 | 0.112907 | 0.016156 | 141,134 | 2 | 2,397 |
| All autosomes (Fig. S2) | 12 | 1,832.3 | 3.9 | 0.192 | 0.116342 | 0.016899 | 145,428 | 2 | 2,580 |
| Also chr. 9 excluded (Fig. S2) | 10 | 1,542.0 | 8.2 | 0.166 | 0.098351 | 0.014296 | 122,939 | 2 | 3,214 |

Callable: sequence passing depth and quality filters. θ₀ and ρ₀: scaled mutation and recombination rates per 100-bp bin. *N*₀ = θ₀/(4μ × 100) with μ = 2 × 10^−9^ per site per generation. Min. recomb.: smallest number of inferred recombination events across the 29 free parameters (Li & Durbin 2011 recommend ≥ 10). PSMC 0.6.5-r67, -N25 -t15 -r5 -p "2+2+25*2+4+6", 100 bootstrap replicates per model. Missing data in the masked models include the masked inversion.

**Table S4:** Habitat codings for *Somatochlora*

**Table S5.** Branches on the crown-to-*S. arctica* path and posterior placement of the most recent peatland gain (primary coding; 1,000 maps).

| **Branch** | **Description** | **Span (Ma)** | **Median gain age (Ma)** | **Maps (%)** |
| --- | --- | --- | --- | --- |
| — | before the crown (specialist throughout) | ≥ 23.7 | — | 15.4 |
| a | stem of 39-sp. clade | 23.7–19.7 | 22.4 | 1.6 |
| b | stem of 18-sp. clade | 19.7–17.8 | 18.6 | 1.6 |
| c | stem of 17-sp. clade | 17.8–15.9 | 16.7 | 3.8 |
| d | stem of 8-sp. clade | 15.9–13.7 | 14.9 | 4.0 |
| e | stem of 5-sp. clade | 13.7–5.7 | 9.4 | 64.4 |
| f | terminal *S. arctica* | 5.7–0.0 | 2.4 | 9.2 |

**Supplemental Note S1: Implementation of the PSMC**

We mapped the reads to their respective reference genomes with minimap2 2.28 map-hifi (Li 2018) and sorted and merged the alignments into BAM files with samtools 1.19.2 (Danecek et al. 2021). We calculated alignment depth with mosdepth 0.3.7 at MAPQ ≥ 20 (Pedersen & Quinlan 2018). We restricted the analyses to chromosome-scale autosomes, excluding unplaced scaffolds and the X chromosomes, which we identified by their half-autosomal depth and near-zero heterozygosity (Table S2). We further excluded sites with depth below 10 or one third of the median autosomal depth (D; whichever was higher) or above 2D (Nadachowska-Brzyska et al. 2016), giving filters of 10–56× for *S. meridionalis* (D = 28.2×) and 16–97× for *S. arctica* (D = 48.5×). We generated diploid consensus sequences with bcftools 1.19 (mpileup -X pacbio-ccs -q20 -Q20 | call -c; Danecek et al. 2021), applied the depth filters with vcfutils.pl vcf2fq (Li et al. 2009), and converted the output to PSMC input format with fq2psmcfa -q20. We ran PSMC 0.6.5-r67 with the parameters -N25 -t15 -r5 -p "2+2+25*2+4+6", which split the first time window to avoid false recent peaks (Hilgers et al. 2025). We ran 100 bootstrap replicates per model and confirmed that every free parameter was supported by at least 10 inferred recombination events (Li & Durbin 2011; Table S3). The curation notes of the *S. arctica* assembly report a heterozygous inversion on chromosome 5 (65.88–152.04 Mb; 86.16 Mb) and a microchromosome (chromosome 12) whose order and orientation are uncertain, and we further found that chromosome 9 had more than twice the median autosomal heterozygosity (Table S2). To test whether these features influenced the inferred demographic history, we repeated the *S. arctica* PSMC twice: once with the microchromosome excluded and the chromosome 5 inversion masked, and once with chromosome 9 also excluded (Fig. S1; Table S3). We scaled the PSMC output assuming a genome-wide mutation rate of μ = 2×10⁻⁹ per site per generation (Liu et al. 2017; Tolman et al. 2025) and generation times of 3 years for *S. meridionalis* and 2 years for *S. arctica*. We plotted iteration 20 of each run and omitted the final merged free parameter (the last six atomic time intervals) from the plots.

**Supplemental Note S2: Ancestral State Reconstruction of peatland Specialization *Somatochlora***

To conduct ancestral state reconstruction, we used the dated tree of *Somatochlora* from Goodman et al. (2025), pruned to *Somatochlora* in ape (Paradis & Schliep 2019). We searched the literature for evidence of whether each sampled *Somatochlora* species is considered a peatland habitat specialist. In our primary coding, 9 species were coded as specialists, 26 as non-specialists, and 5 with a flat tip prior (P = 0.5 per state). The flat-prior species were *S. calverti*, *S. daviesi*, *S. borisi*, and *S. shanxiensis*, for which evidence of habitat use is not available, and *S. forcipata*, which is strongly peatland-associated but may also use streams and swamps (Table S4). We fit continuous-time Markov (Mk) models for discrete characters (Pagel 1994; Lewis 2001) by maximum likelihood with fitMk in phytools (Revell 2012, 2024). We compared equal-rates (ER) and all-rates-different (ARD) models, in which gain and loss rates are equal or estimated separately, respectively, and specified root-state probabilities as proportional to their conditional likelihoods (FitzJohn et al. 2009). We assessed model support with small-sample corrected AIC (AICc; Hurvich & Tsai 1989) converted to Akaike weights (Burnham & Anderson 2002). We estimated marginal ancestral-state probabilities at internal nodes under both models with ancr (phytools) and averaged them across models by Akaike weight.

To estimate the number of gains and losses of peatland specialization across the tree, and when the *S. arctica* lineage most recently gained this state, we used stochastic character maps (Nielsen 2002; Huelsenbeck et al. 2003; Bollback 2006) simulated with make.simmap (phytools), conditioning on each model's maximum-likelihood rate matrix and the same root prior. We simulated 1,000 maps per analysis, allocated between ER and ARD in proportion to their Akaike weights (primary coding: ER = 543, ARD = 457), and summarized the posterior number of gains and losses and the posterior mean proportion of each branch spent in the peatland specialist state. For each map, we identified the most recent gain of peatland specialization on the lineage leading to *S. arctica*, i.e. the start of the uninterrupted run of specialization ending at *S. arctica*; earlier gains that were later lost were ignored. Maps in which the whole crown-to-tip path was in the peatland state were classified as 'specialist since the crown' (gain at or before the crown age). We summarized the timing of this gain as the median and 95% highest-density interval (HDI; Hyndman 1996). Analyses were conducted in R 4.4.3 (R Core Team 2025); trees were read with treeio (Wang et al. 2020), and figures were produced with ggtree (Yu et al. 2017), ggplot2 (Wickham 2016), ggtext (Wilke & Wiernik 2022), and patchwork (Pedersen 2024).

To assess the sensitivity of our ancestral state reconstructions to habitat coding, we repeated this procedure under three alternative codings, each modifying the primary coding: 1) *S. forcipata* coded as a specialist (instead of unknown); 2) *S. forcipata*, *S. kennedyi*, *S. sahlbergi*, *S. walshii*, and *S. flavomaculata* all coded as specialists; and 3) the two qualified specialist codings (*S. alpestris* and *S. margarita*; Table S4) recoded as non-specialists.

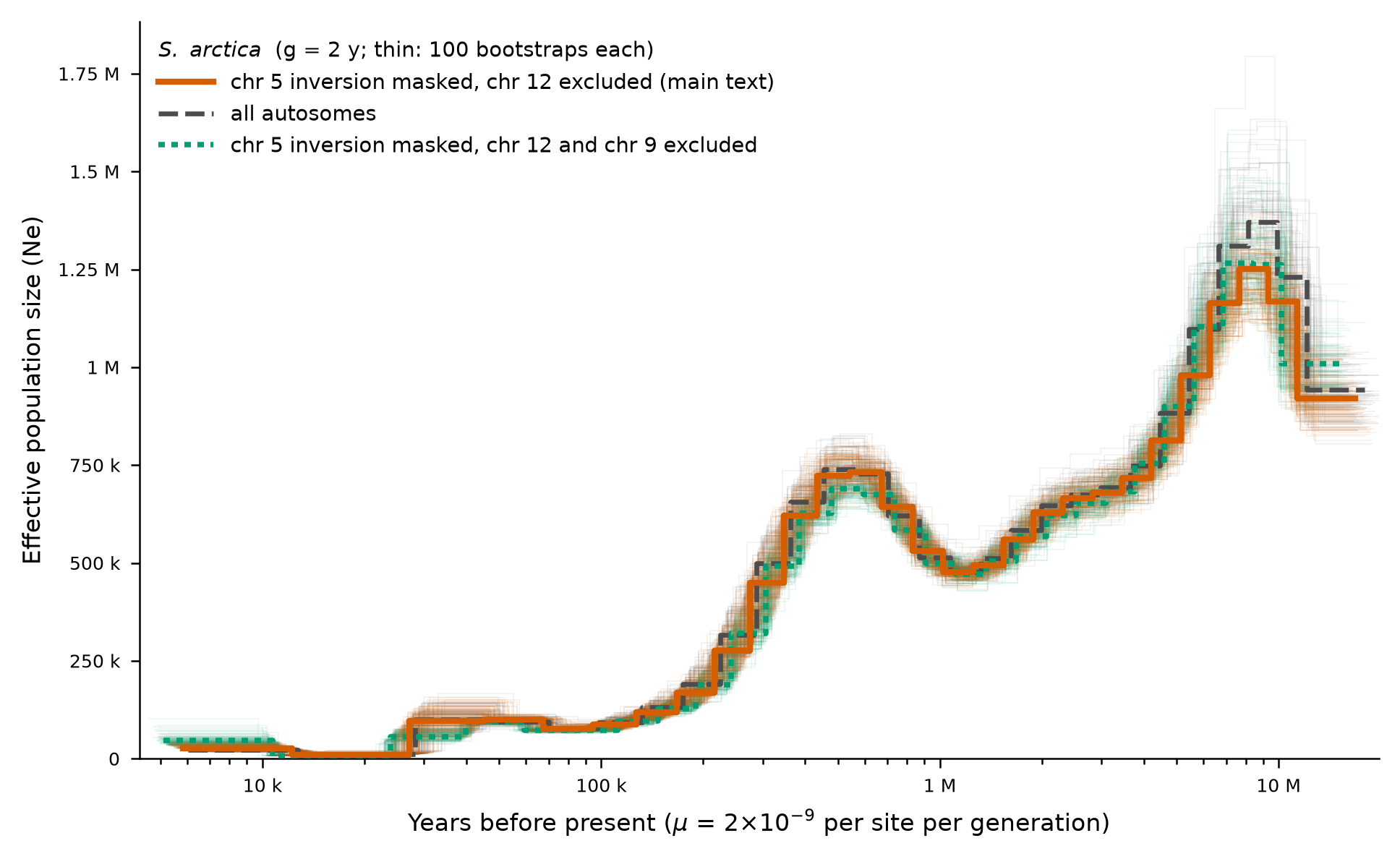

**Figure S1.** Sensitivity of the *Somatochlora arctica* PSMC to genome features. Effective population size (N_e_) inferred with the chromosome 5 inversion masked and the microchromosome (chromosome 12) excluded (solid orange; model shown in Fig. 1A), with all autosomes (dashed gray), and with chromosome 9 additionally excluded (dotted green). Thin lines show 100 bootstrap replicates per model. Scaled with μ = 2×10⁻⁹ per site per generation and g = 2 years.

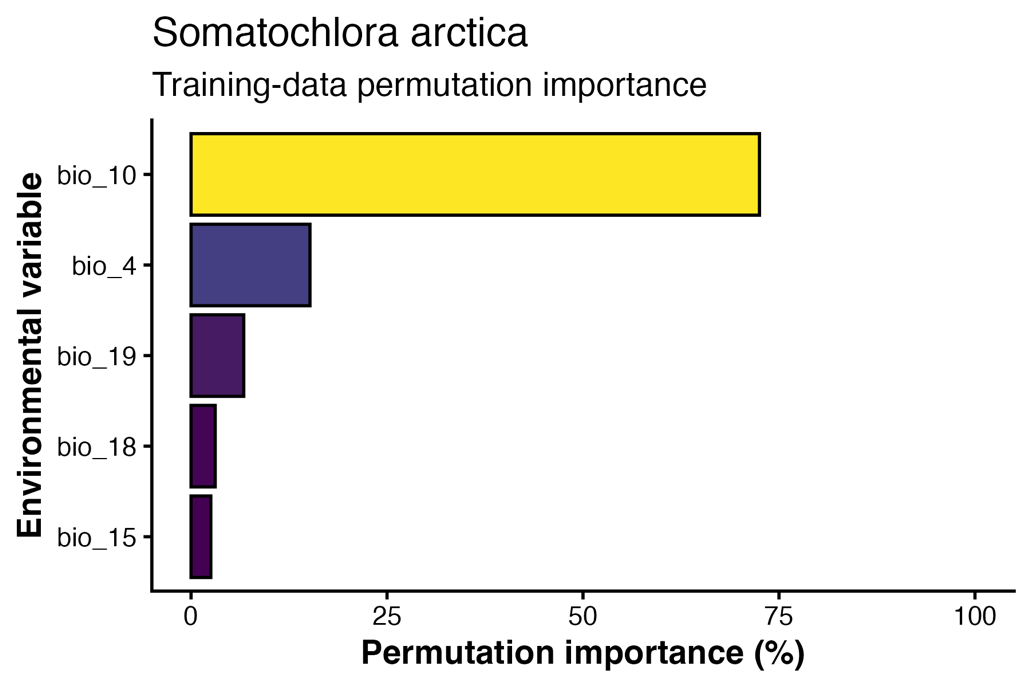

**Figure S2.** Training-data permutation importance of environmental predictors in the optimal Maxent model for *Somatochlora arctica*. Importance was calculated by randomly permuting each predictor among presence and background points and measuring the resulting decrease in training AUC. Positive decreases were normalized to percentages; longer bars indicate greater model dependence on that predictor. Variables are mean temperature of the warmest quarter (bio10), temperature seasonality (bio04), precipitation seasonality (bio15), precipitation of the warmest quarter (bio18), and precipitation of the coldest quarter (bio19)

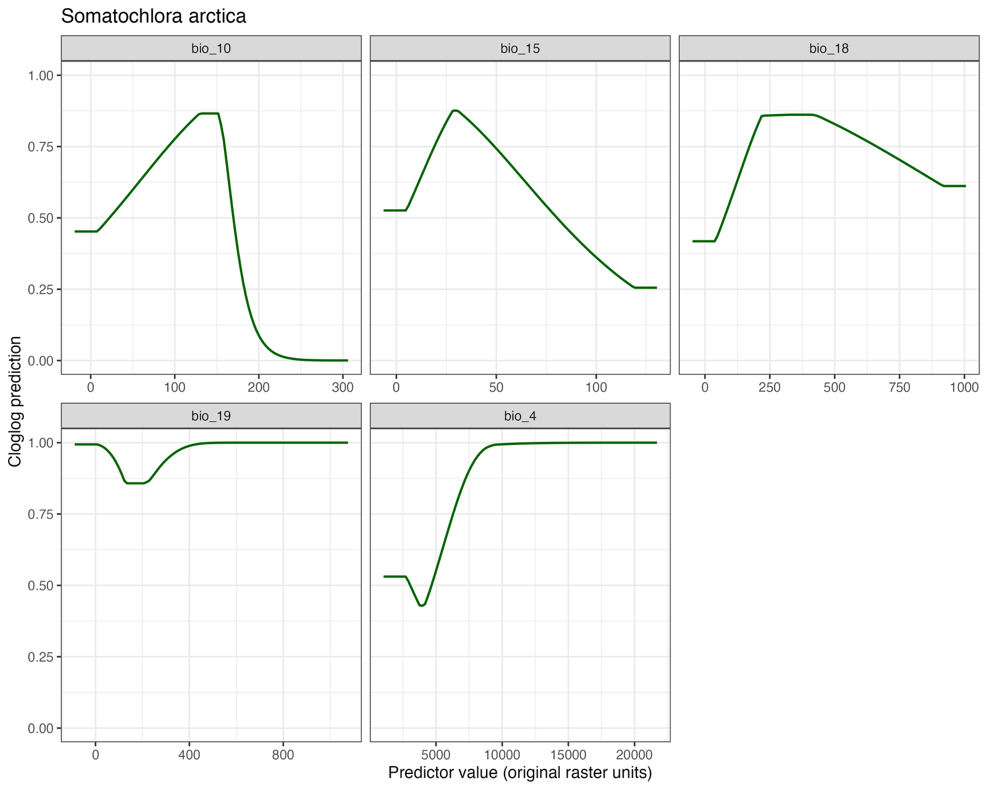

**Figure S3.** Conditional response curves of environmental predictors in the optimal Maxent model for *Somatochlora arctica*. Each curve shows the relationship between one predictor and cloglog suitability while all other predictors are held at their model sample means. Variables are mean temperature of the warmest quarter (bio10), temperature seasonality (bio04), precipitation seasonality (bio15), precipitation of the warmest quarter (bio18), and precipitation of the coldest quarter (bio19). Predictor values are shown in original raster units, with bio10 expressed as degrees Celsius multiplied by 10; bio15 is the coefficient of variation (%), and bio18 and bio19 are in millimeters.

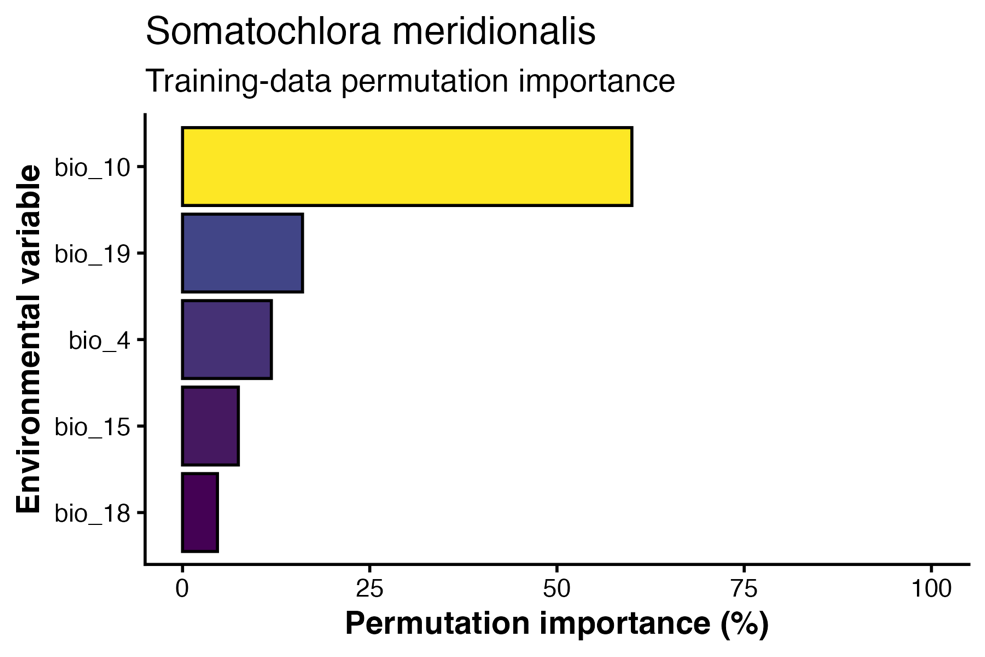

**Figure S4.** Training-data permutation importance of environmental predictors in the optimal Maxent model for *Somatochlora meridionalis*. Importance was calculated by randomly permuting each predictor among presence and background points and measuring the resulting decrease in training AUC. Positive decreases were normalized to percentages; longer bars indicate greater model dependence on that predictor. Variables are mean temperature of the warmest quarter (bio10), temperature seasonality (bio04), precipitation seasonality (bio15), precipitation of the warmest quarter (bio18), and precipitation of the coldest quarter (bio19).

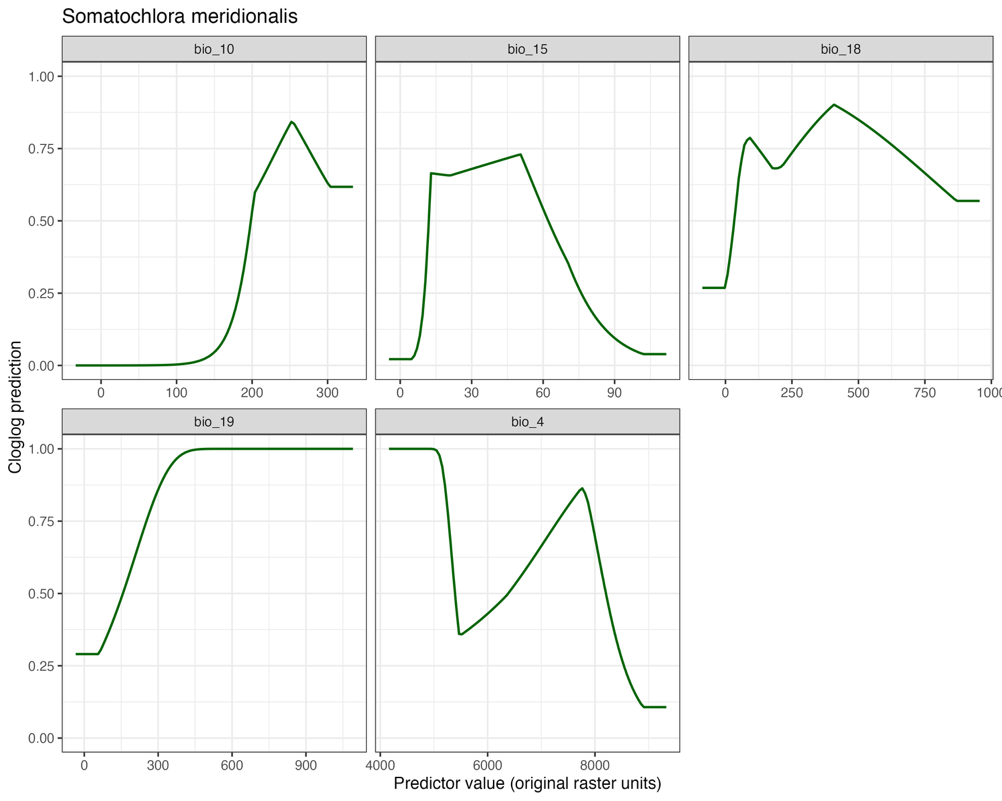

**Figure S5.** Conditional response curves of environmental predictors in the optimal Maxent model for *Somatochlora meridionalis*. Each curve shows the relationship between one predictor and cloglog suitability while all other predictors are held at their model sample means. Variables are mean temperature of the warmest quarter (bio10), temperature seasonality (bio04), precipitation seasonality (bio15), precipitation of the warmest quarter (bio18), and precipitation of the coldest quarter (bio19). Predictor values are shown in original raster units, with bio10 expressed as degrees Celsius multiplied by 10; bio15 is the coefficient of variation (%), and bio18 and bio19 are in millimeters.

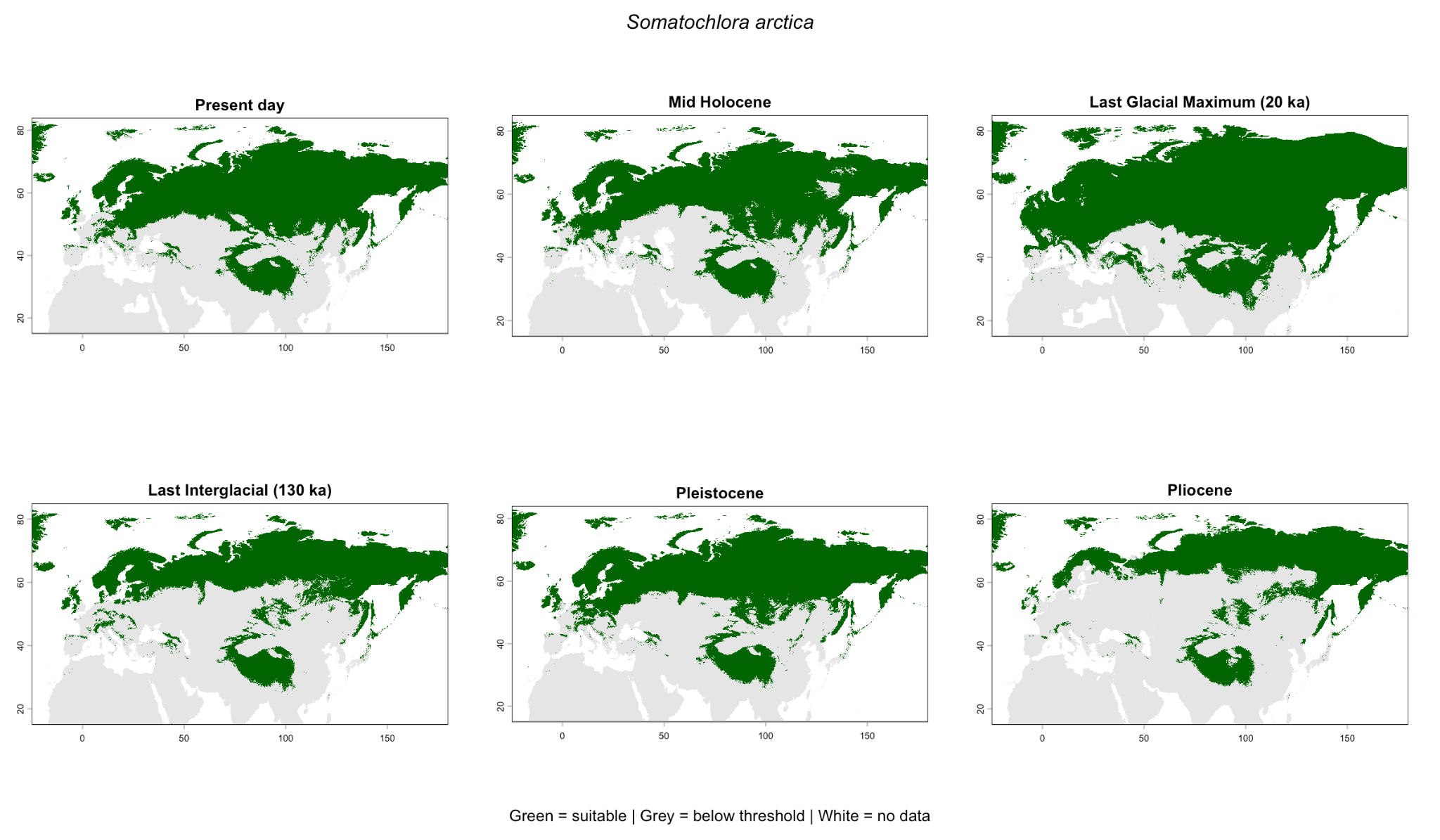

**Figure S6.** Binary Maxent predictions of climatic suitability for *Somatochlora arctica* across the Palearctic. Panels show, from left to right, the present, mid-Holocene, and Last Glacial Maximum (top row), and Last Interglacial, Pleistocene MIS19, and Pliocene MIS M2 (bottom row). Predictions were generated from the optimal model and classified using the 10th percentile of present-day suitability at retained occurrence localities, with the same threshold applied to all paleoclimate projections. Green indicates suitable climate, gray indicates values below the threshold, and white indicates no data. Predictions are shown before application of the independent ice masks used for suitable-area calculations.

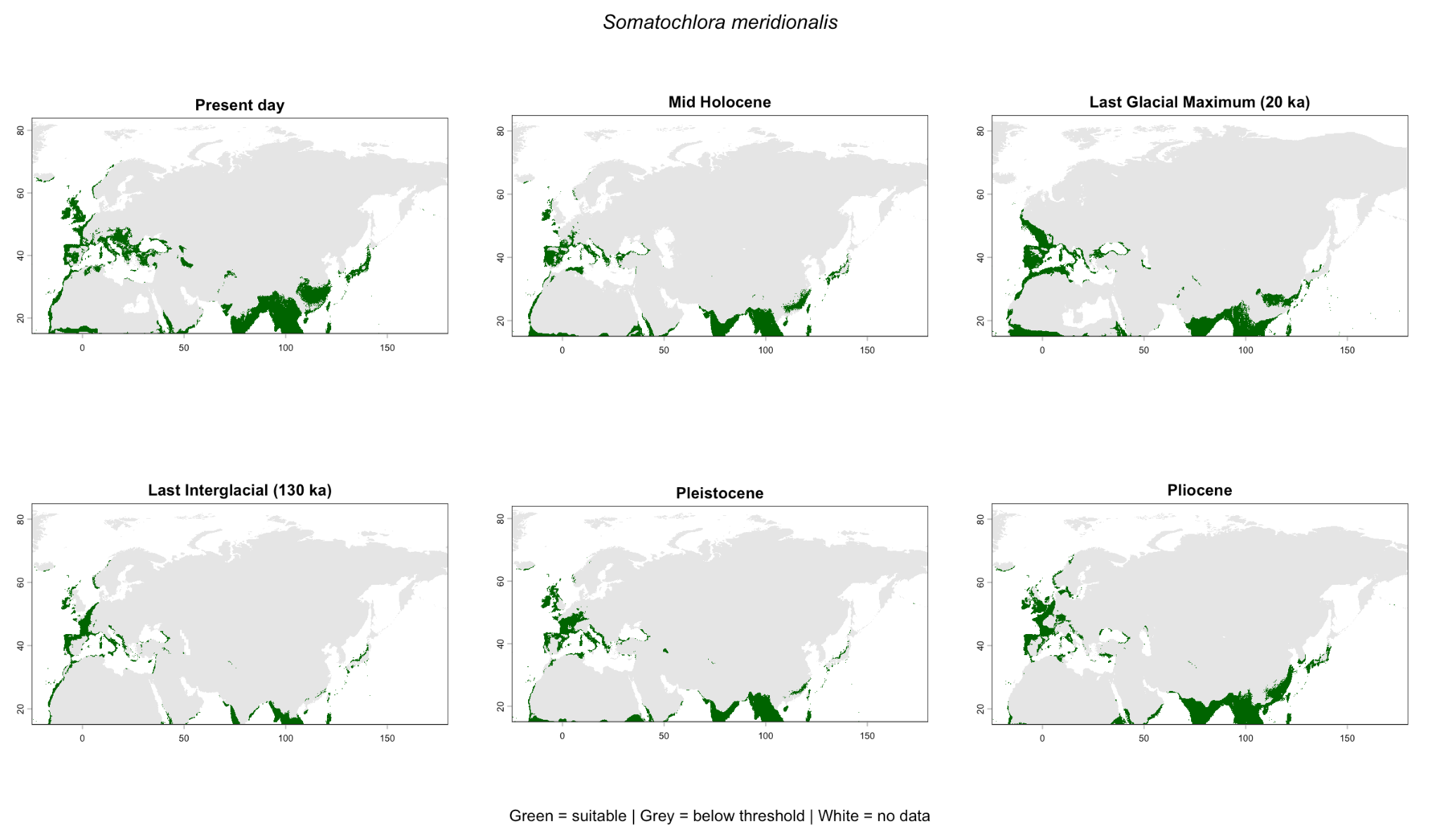

**Figure S7.** Binary Maxent predictions of climatic suitability for *Somatochlora meridionalis* across the Palearctic. Panels show, from left to right, the present, mid-Holocene, and Last Glacial Maximum (top row), and Last Interglacial, Pleistocene MIS19, and Pliocene MIS M2 (bottom row). Predictions were generated from the optimal model and classified using the 10th percentile of present-day suitability at retained occurrence localities, with the same threshold applied to all paleoclimate projections. Green indicates suitable climate, gray indicates values below the threshold, and white indicates no data. Predictions are shown before application of the independent ice masks used for suitable-area calculations.

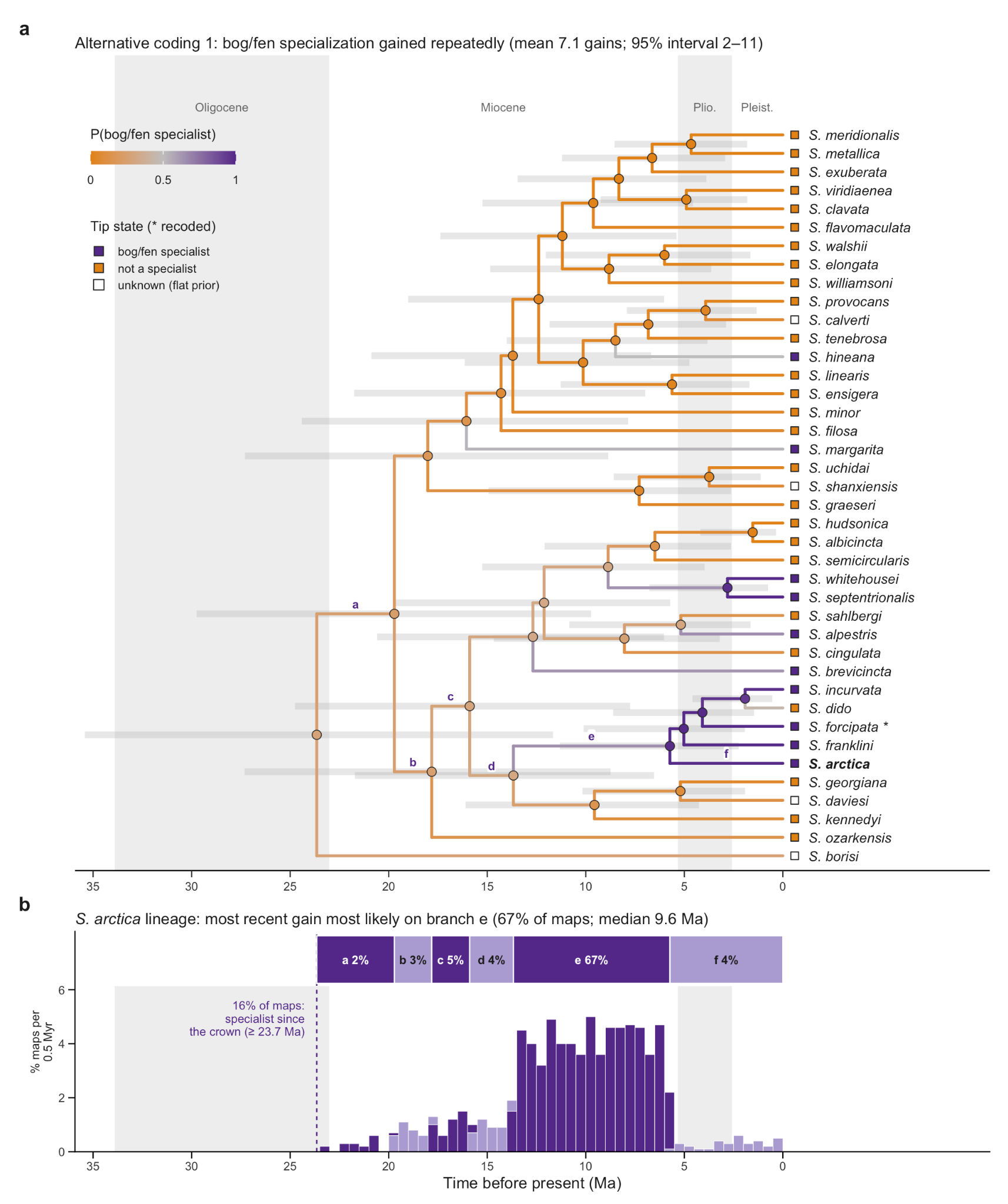

**Figure S8.** Ancestral state reconstruction of peatland specialization in *Somatochlora* under alternative coding 1, with *S. forcipata* coded as a peatland specialist. Tips whose coding differs from the primary analysis are marked with an asterisk. (a) Tree and ancestral states as in Fig. 2a. (b) Posterior age of the most recent peatland gain on the *S. arctica* lineage as in Fig. 2b.

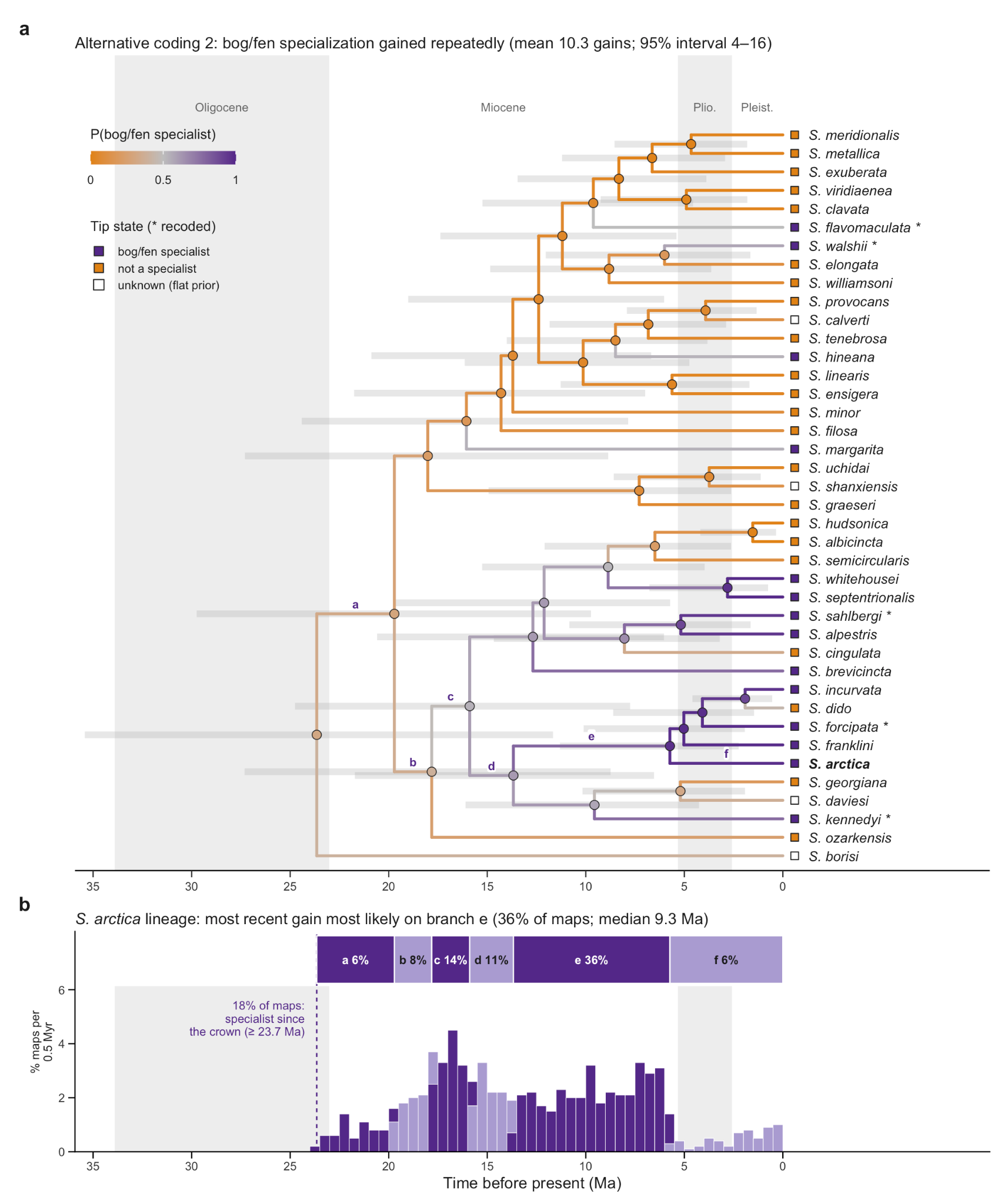

**Figure S9.** Ancestral state reconstruction of peatland specialization in *Somatochlora* under alternative coding 2, with *S. forcipata*, *S. kennedyi*, *S. sahlbergi*, *S. walshii*, and *S. flavomaculata* coded as specialists. Tips whose coding differs from the primary analysis are marked with an asterisk. (a) Tree and ancestral states as in Fig. 2a. (b) Posterior age of the most recent peatland gain on the *S. arctica* lineage as in Fig. 2b.

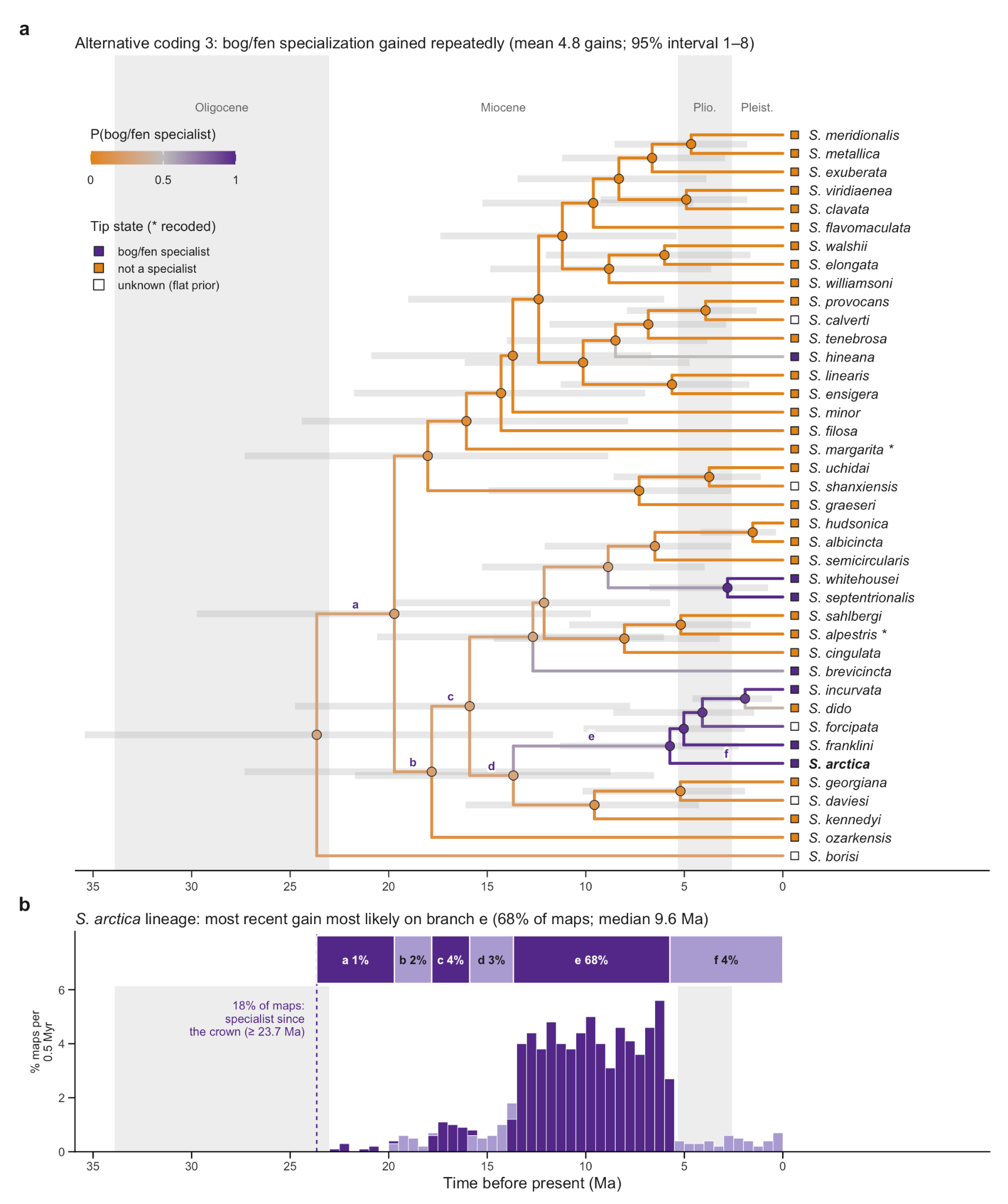

**Figure S10.** Ancestral state reconstruction of peatland specialization in *Somatochlora* under alternative coding 3, with *S. alpestris* and *S. margarita* recoded as non-specialists (*S. forcipata* unknown). Tips whose coding differs from the primary analysis are marked with an asterisk. (a) Tree and ancestral states as in Fig. 2a. (b) Posterior age of the most recent peatland gain on the *S. arctica* lineage as in Fig. 2b.
